# Abscisic acid is essential for rewiring of jasmonic acid-dependent defenses during herbivory

**DOI:** 10.1101/747345

**Authors:** Irene A Vos, Adriaan Verhage, Lewis G Watt, Ido Vlaardingerbroek, Robert C Schuurink, Corné MJ Pieterse, Saskia CM Van Wees

## Abstract

Jasmonic acid (JA) is an important plant hormone in the regulation of defenses against chewing herbivores and necrotrophic pathogens. In *Arabidopsis thaliana*, the JA response pathway consists of two antagonistic branches that are regulated by MYC- and ERF-type transcription factors, respectively. The role of abscisic acid (ABA) and ethylene (ET) in the molecular regulation of the MYC/ERF antagonism during plant-insect interactions is still unclear. Here, we show that production of ABA induced in response to leaf-chewing *Pieris rapae* caterpillars is required for both the activation of the MYC-branch and the suppression of the ERF-branch during herbivory. Exogenous application of ABA suppressed ectopic ERF-mediated *PDF1.2* expression in *35S::ORA59* plants. Moreover, the GCC-box promoter motif, which is required for JA/ET-induced activation of the ERF-branch genes *ORA59* and *PDF1.2*, was targeted by ABA. Application of gaseous ET counteracted activation of the MYC-branch and repression of the ERF-branch by *P. rapae*, but infection with the ET-inducing necrotrophic pathogen *Botrytis cinerea* did not. Accordingly, *P. rapae* performed equally well on *B. cinerea*-infected and control plants, whereas activation of the MYC-branch resulted in reduced caterpillar performance. Together, these data indicate that upon feeding by *P. rapae*, ABA is essential for activating the MYC-branch and suppressing the ERF-branch of the JA pathway, which maximizes defense against caterpillars.

## Introduction

In nature plants are a food source for over one million herbivorous insect species (Howe and Jander, 2008). The evolutionary arms race between plants and their herbivorous insect enemies has led to a highly sophisticated defense system in plants that can recognize wounding and oral secretion of the insects and respond with the production of nutritive value-diminishing enzymes, toxic compounds, or predator-attracting volatiles (Kessler and Baldwin, 2002; Lawrence and Koundal, 2002; Wittstock et al., 2004; Chen et al., 2005; Mithöfer and Boland, 2012; Dicke, 2016). Conversely, insects can estimate the quality and suitability of the plant as a food source by contact chemoreceptors on the insect mouthparts, antennae and tarsi (Howe and Jander, 2008; Appel and Cocroft, 2014; Dicke, 2016). Because plant defenses are costly, they are often only activated in case of insect or pathogen attack and not constitutively expressed (Walters and Heil, 2007; Vos et al., 2013a). The induced immune response is shaped by the induced production of diverse plant hormones. The quantity, composition and timing of the hormonal blend tailors the defense response specifically to the attacker at hand, thereby prioritizing effective over ineffective defenses and minimizing fitness costs (De Vos et al., 2005; Pieterse et al., 2012; Vos et al., 2013a; Vos et al., 2015).

Infestation with chewing herbivores or infection with necrotrophic pathogens triggers the production of the plant hormone jasmonic acid (JA), and its bioactive derivative JA-Ile (Creelman et al., 1992; Penninckx et al., 1996). Binding of JA-Ile to the JA receptor complex consisting of the F-box protein COI1 and a JAZ repressor protein (Xie et al., 1998; Yan et al., 2009; Sheard et al., 2010), leads to degradation of JAZ proteins via the 26S proteasome pathway (Chini et al., 2007; Thines et al., 2007). Without JA, JAZ proteins repress JA-responsive gene expression by binding to transcriptional activators, such as MYC2, EIN3 and EIL1 (Pauwels and Goossens, 2011; Song et al., 2014b; Caarls et al., 2015). When JA accumulates the JAZ proteins are degraded thereby releasing transcription factors that can activate JA-regulated genes.

Within the JA pathway, two distinct, antagonistic branches of transcriptional regulation are recognized; the MYC-branch and the ERF-branch. Feeding by chewing herbivores activates the MYC-branch (Verhage et al., 2011; Vos et al., 2013b). This branch is controlled by the basic helix-loop-helix leucine zipper transcription factors MYC2, MYC3 and MYC4 leading to transcription of hundreds of JA-responsive MYC-branch regulated genes, including *VSP1* and *VSP2* (Anderson et al., 2004; Lorenzo et al., 2004; Fernández-Calvo et al., 2011; Niu et al., 2011). Furthermore, previous studies have indicated that ABA plays a co-regulating role in the activation of the MYC-branch (Anderson et al., 2004; Bodenhausen and Reymond, 2007; Sánchez-Vallet et al., 2012; Vos et al., 2013b). For example, in the ABA-deficient mutant *aba2-1*, expression of the JA-responsive gene *VSP1* was reduced upon feeding by caterpillars of *Pieris rapae* (small cabbage white) compared to wild-type Col-0 plants (Vos et al., 2013b). In contrast to the herbivore-induced MYC-branch, the ERF-branch is activated upon infection with necrotrophic pathogens. The transcription factors EIN3 and EIL1 and the ERF transcription factors ERF1 and ORA59 activate a large set of JA-responsive ERF-branch regulated genes, including *PDF1.2* (Caarls et al., 2015). The expression of *ERF1*, *ORA59* and *PDF1.2* is impaired in both JA- and ethylene (ET)-insensitive mutants, indicating that joint activation of the JA and ET pathways is necessary for full expression of the ERF-branch (Penninckx et al., 1998; Lorenzo et al., 2003; Pré et al., 2008; Broekgaarden et al., 2015).

It has been shown that the ABA co-regulated MYC-branch and the ET co-regulated ERF-branch of the JA pathway antagonize each other. For example, upon infestation with *P. rapae* caterpillars, the MYC-branch is activated, while the ERF-branch is suppressed (Verhage et al., 2011; Vos et al., 2013b). In *myc2* mutant plants, *ORA59* and *PDF1.2* expression was highly upregulated after feeding by *P. rapae*, indicating that in wild-type plants, MYC2 represses *ORA59* and *PDF1.2* expression after feeding by *P. rapae* (Verhage et al., 2011; Vos et al., 2013b). Additionally, exogenously applied ABA had a positive effect on expression of the MYC-branch after feeding by *P. rapae* (Vos et al., 2013b) and caused suppression of *PDF1.*2 induction after exogenous application of JA (Anderson et al., 2004). Recently, it was shown that the MYC-branch transcription factors MYC2, MYC3 and MYC4 interact with the ERF-branch transcription factors EIN3 and EIL1 and that they repress each other’s transcriptional activity (Song et al., 2014a).

These antagonistic effects between the MYC- and ERF-branch on gene expression levels also have an effect on plant resistance. ABA-deficient mutants have been reported to be more susceptible to herbivory (Thaler and Bostock, 2004; Bodenhausen and Reymond, 2007; Dinh et al., 2013) and more resistant to necrotrophic pathogens (Anderson et al., 2004; Sánchez-Vallet et al., 2012). Conversely, ET insensitive mutants are in general more susceptible to necrotrophic pathogens and more resistant to herbivorous insects compared to wild-type plants (Van Loon et al., 2006; Broekgaarden et al., 2015). Hence, the interplay between the MYC- and the ERF-branch may allow the plant to activate a specific set of JA-responsive genes that is required for an optimal defense against the attacker encountered (Pieterse et al., 2012).

To study the role of ABA and ET in the molecular regulation of the MYC/ERF balance in *Arabidopsis thaliana* (hereafter Arabidopsis) upon attack by *P. rapae*, we analyzed hormone signaling mutants for their gene expression response, hormone production and defense against *P. rapae*. We provide evidence that after *P. rapae* infestation ABA accumulation plays an essential modulating role in the activation of the MYC-branch, possibly by activating the MYC2, MYC3 and MYC4 transcription factors. Concomitantly, ABA can suppress the ERF-branch independently of the MYC transcription factors, by targeting the GCC-box, which is present in the promoters of *ORA59* and *PDF1.2*. Furthermore, activation of the MYC-branch, either by application of JA or ABA or by using the *ein2-1* mutant, resulted in a negative effect on caterpillar performance, whereas activation of the ERF-branch by infection with the necrotrophic pathogen *Botrytis cinerea* did not.

## Results

### ABA- and ET-dependency of JA-dependent defense gene expression upon *P. rapae* feeding

The JA-dependent transcriptional response of Arabidopsis to *P. rapae* feeding is predominantly regulated through activation of the MYC-branch of the JA pathway and concomitant suppression of the ERF-branch (Verhage et al., 2011). Here, we investigated whether ABA and ET have a role in the differential expression of the MYC- and the ERF-branch during induction of JA-dependent defense signaling by *P. rapae* feeding. Expression of the MYC-branch marker gene *VSP2* and the ERF-branch marker gene *PDF1.2* was monitored in wild-type Col-0, *MYC2*-impaired mutant *jin1-7* (hereafter called *myc2*), *MYC2*, *MYC3*, *MYC4* triple mutant *myc2,3,4*, ABA biosynthesis mutant *aba2-1* and ET response mutant *ein2-1*. First-instar *P. rapae* caterpillars were allowed to feed for 24 h on the different Arabidopsis genotypes, after which they were removed. Comparable to Col-0, *ein2-1* plants showed strong *P. rapae-*induced transcription of *VSP2* at 24 h and 30 h (Figure 1). *VSP2* transcript levels decreased to basal levels at 48 h in both Col-0 and *ein2-1* plants, suggesting that stimulation of the MYC-branch lasted until at least 6 h after removal of the caterpillars. At 24 h, the *ein2-1* plants showed a significantly enhanced transcription level of *VSP2* compared with Col-0 (Figure 1), indicating a primed responsiveness to the MYC-branch. *PDF1.2* transcript levels were very low in both Col-0 and *ein2-1*. In *myc2* as well as in *aba2-1* mutants, the transcriptional patterns of *VSP2* and *PDF1.2* were opposite to those observed in Col-0, showing low *VSP2* expression and high *PDF1.2* expression up to 30 h. In *myc2,3,4* mutants, expression of *VSP2* was almost zero. *PDF1.2* levels in *myc2,3,4* plants were similar to Col-0 up to 30 h, after which they increased significantly at 48 h (Figure 1). Together these results confirm that the MYC transcription factors function as a switch between the two branches of the JA pathway, whereby *myc2,3,4* plants show a delay in expression of the ERF-branch. Furthermore, ABA is essential for activation of the MYC-branch and repression of the ERF-branch upon *P. rapae* feeding, while the ET pathway has only a small, though significant, effect on the MYC/ERF-balance during *P. rapae* feeding.

**Figure 1:**
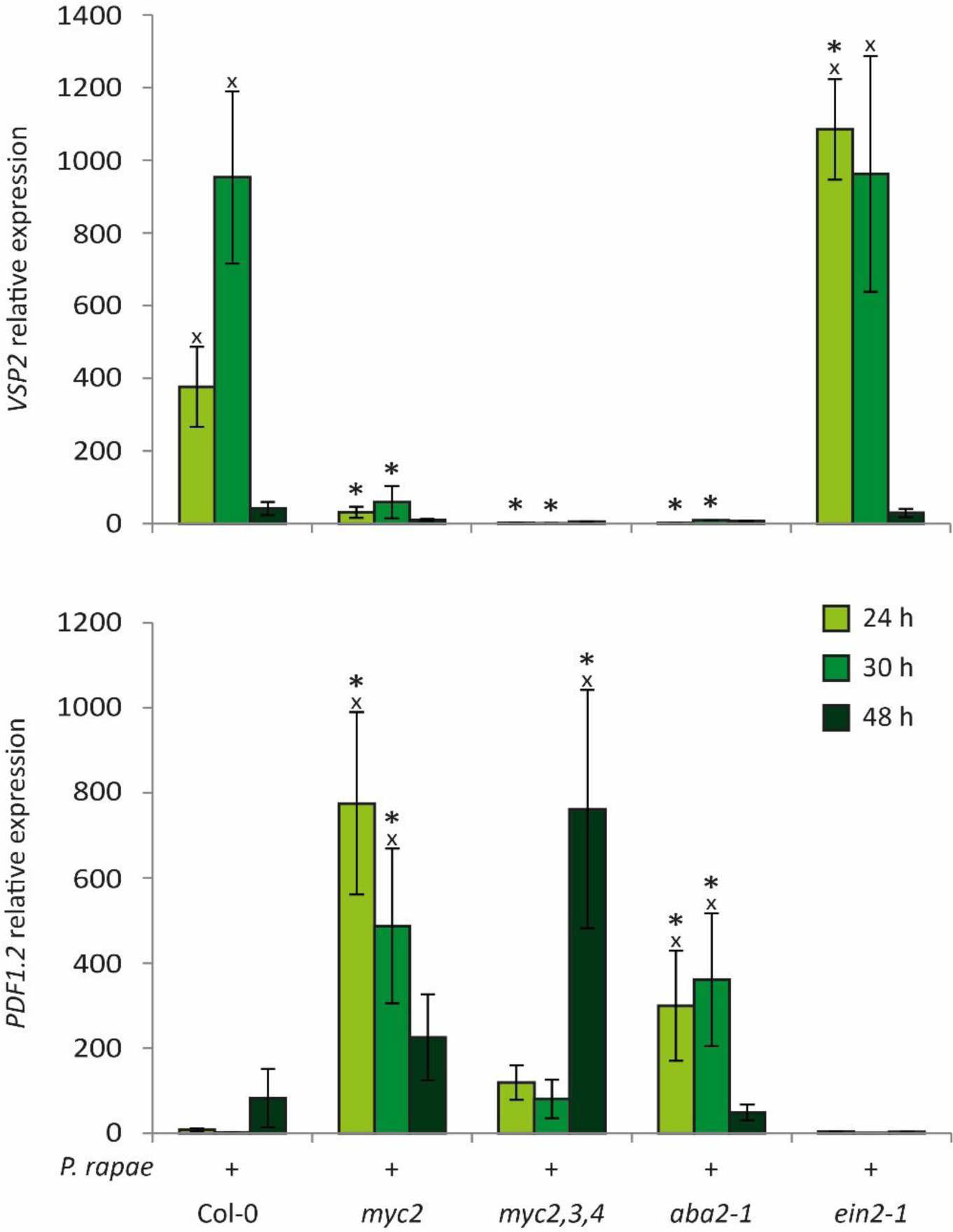
Expression of JA-responsive MYC- and ERF-branch marker genes in response to *P. rapae* feeding in Arabidopsis mutants. RT-qPCR analysis of *VSP2* and *PDF1.2* gene expression in *P. rapae*-infested leaves of Col-0, *myc2*, *myc2,3,4*, *aba2-1* and *ein2-1* plants. Indicated are expression values relative to non-infested Col-0 plants at 24 h after infestation. First-instar *P. rapae* caterpillars were allowed to feed for 24 h after which they were removed. Infested leaves were harvested at the indicated time points after introduction of the caterpillars. Crosses indicate a statistically significant difference with the corresponding non-infested control (expression data of non-infested controls are not shown; two-way ANOVA (treatment x time point), LSD test for multiple comparisons; *P*<0.05). Asterisks indicate a statistically significant difference with Col-0 at the same time point (two-way ANOVA (time point x genotype), LSD test for multiple comparisons; *P*<0.05). Error bars represent SE, *n*=3 plants.

### Hormone accumulation upon *P. rapae* feeding

To study whether the mutants used in this study are affected in herbivore-induced levels of jasmonates (JAs; JA, the biologically highly active conjugate JA-Ile and the JA-precursor OPDA) and ABA we monitored their accumulation in response to *P. rapae* feeding. We also determined the production of ET in Col-0 wild-type plants. For the measurement of JAs and ABA, first-instar caterpillars were allowed to feed for 24 h after which they were removed from the leaves. Subsequently, hormone levels were measured in caterpillar-damaged leaves at different time points after caterpillar removal. Figure 2 shows that *P. rapae* feeding induced the accumulation of JA, JA-Ile, OPDA and ABA in Col-0 wild-type plants, confirming previous findings (Vos et al., 2013b). In *ein2-1* plants OPDA levels increased to a similar extent as in Col-0, but in contrast, enhanced levels of JA-Ile, JA and ABA were detected. This correlates with the observed enhanced *VSP2* expression in *ein2-1* plants upon *P. rapae* feeding (Figure 1). In *myc2* and *aba2-1* plants, the levels of the JAs raised in general to a similar extent as in Col-0 plants, only the JA and JA-Ile levels did not drop to basal levels at 48 h in *aba2-1* (Figure 2). This indicates that the biosynthesis of JAs is not significantly affected by the *myc2* mutation and only relatively late affected by the *aba2-1* mutation. In *myc2,3,4* plants, OPDA levels were significantly reduced at all time points after caterpillar feeding compared with Col-0. JA levels were also reduced in *myc2,3,4*, but only at 24 h. On the contrary, JA-Ile levels were significantly enhanced in *myc2,3,4* plants at 30 h and 48 h compared with Col-0. This suggests that the JA biosynthesis pathway is perturbed in the *myc2,3,4* plants, resulting in low levels of OPDA, but enhanced production of JA-Ile. ABA levels were highly induced by *P. rapae* feeding in Col-0 at 24 h, but not in *myc2* and *myc2,3,4* plants. At later time points the ABA levels dropped in Col-0 and the differences between the *myc* mutants and Col-0 were no longer significant. These data suggest that herbivore-induced ABA biosynthesis is regulated via MYC transcription factors.

**Figure 2:**
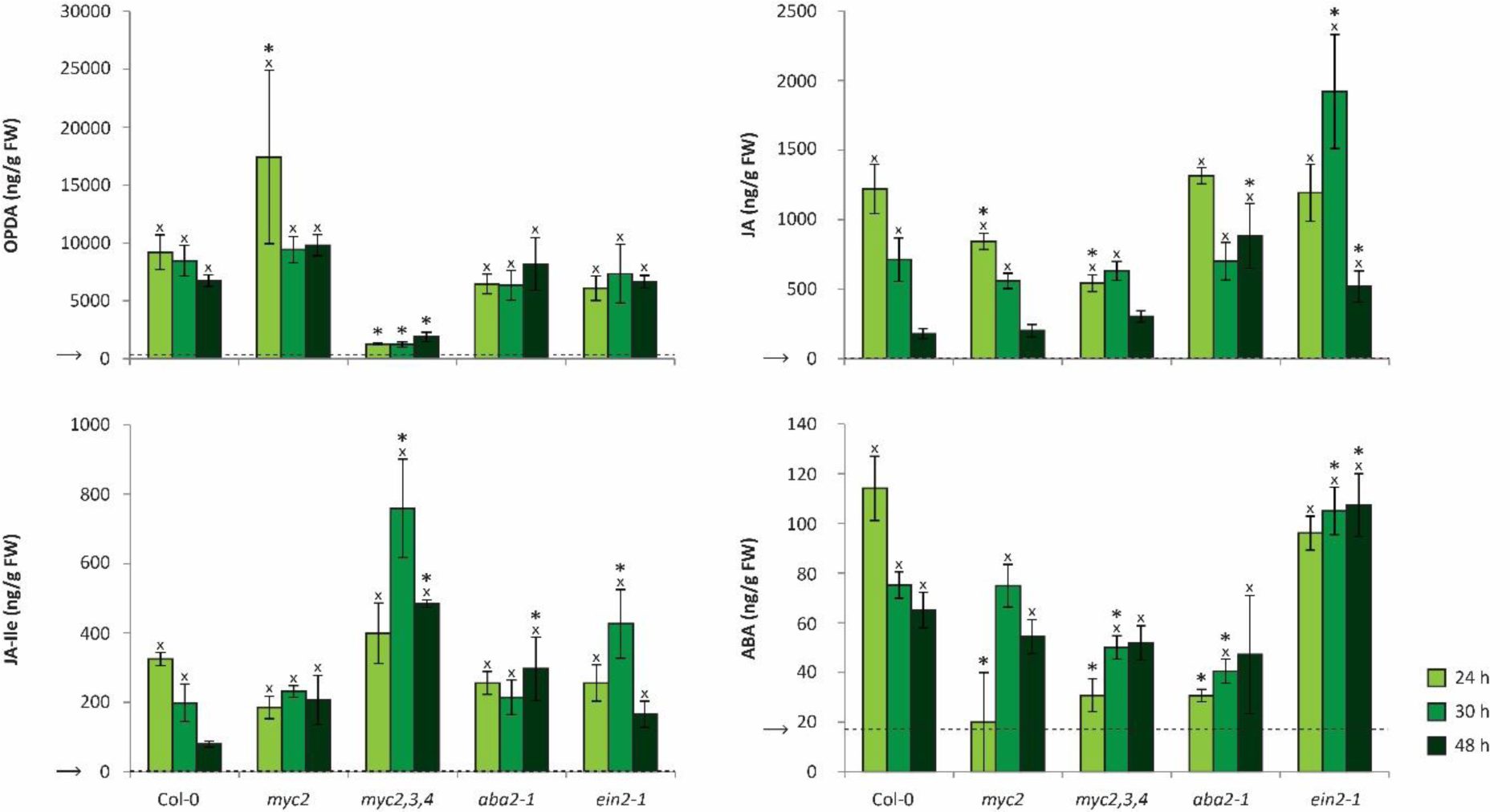
Differential production of OPDA, JA, JA-Ile and ABA in *P. rapae*-infested Arabidopsis mutants. Absolute values (ng/ml/mg FW) of OPDA, JA, JA-Ile and ABA levels that were measured by Triple Quad LC/MS/MS in Col-0, *myc2*, *myc2,3,4*, *aba2-1* and *ein2-1* plants. First-instar *P. rapae* caterpillars were allowed to feed for 24 h after which hormone levels were determined in leaves of non-infested control plants and caterpillar-damaged leaves. Arrows and horizontal dashed lines indicate the average values of non-infested control plants. Crosses indicate a statistically significant difference with the non-infested control of the same line (per genotype two-way ANOVA (treatment x time point), LSD test for multiple comparisons; *P*<0.05). Asterisks indicate a statistically significant difference with Col-0 at the same time point (two-way ANOVA (time point x genotype), LSD test for multiple comparisons; *P*<0.05). Error bars represent SE, *n*=4 plants.

To monitor the emission of ET during *P. rapae* feeding, caterpillar-infested Col-0 plants were placed in 2-l air-tight cuvettes, which allows for continuous ET measurements under climate chamber growth conditions. The positive control, infection with the necrotrophic fungus *B. cinerea,* showed strongly enhanced ET production (Figure 3A), whereas *P. rapae* infestation did not lead to changes in ET production over a 72-h feeding period compared to non-treated control plants (Figure 3B). This indicates that *P. rapae* feeding does not influence ET production.

**Figure 3:**
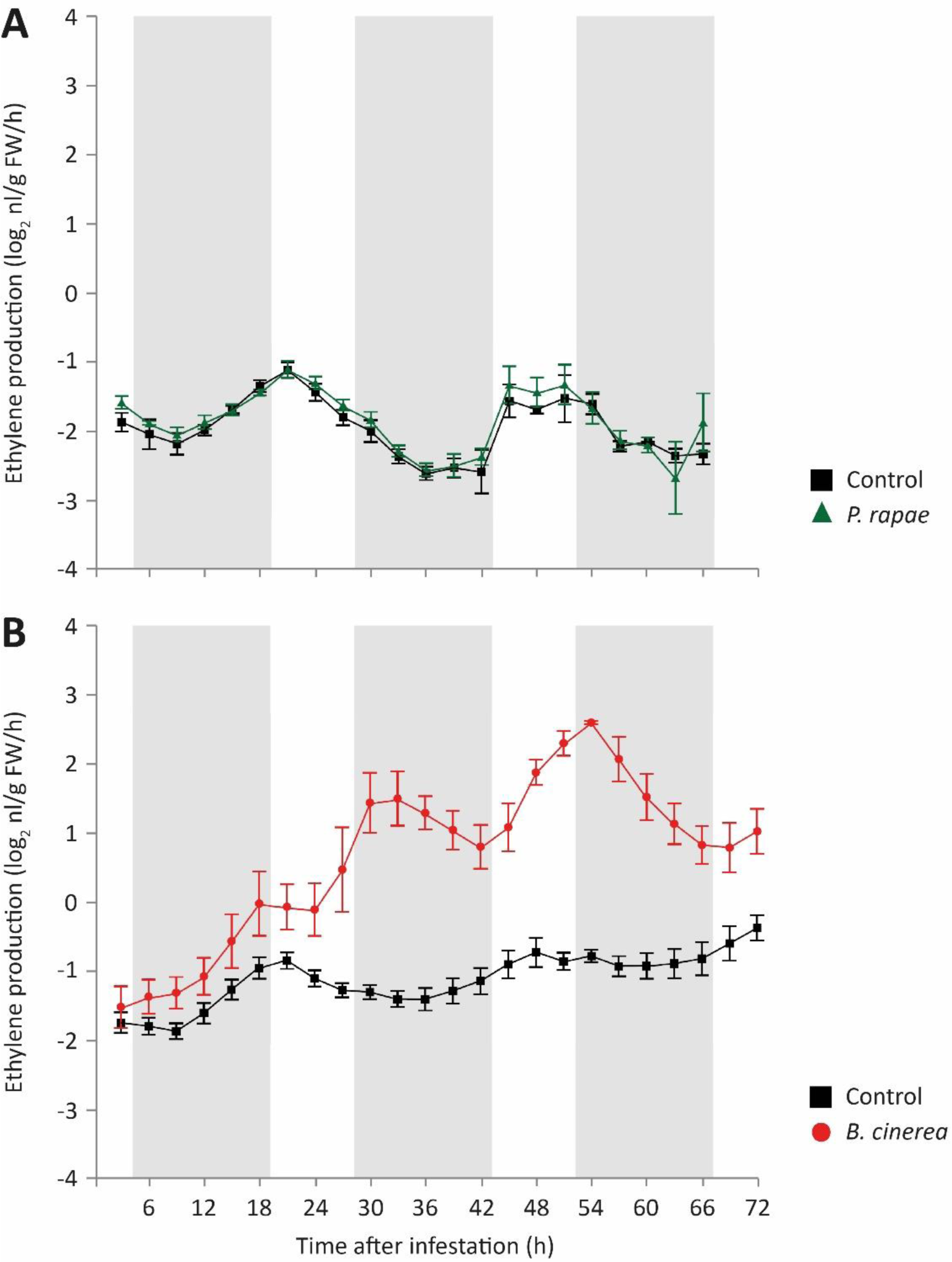
ET production of Col-0 plants during herbivory by *P. rapae* and infection with *B. cinerea*. ET production was monitored in consecutive 3-h time intervals. Col-0 plants were infected with *B. cinerea* (A) or infested with first-instar *P. rapae* caterpillars (B; caterpillars fed on the leaves for the duration of the experiment) in 2-l air-tight cuvettes that were connected to a photoacoustic detection system, which allowed continuous detection of ET levels in the flush-through airflow. Error bars represent SE, *n=*6 plants. White areas indicate the light period, shaded areas indicate the dark period.

### The role of ABA in regulation of MYC/ERF antagonism

To further investigate the role of ABA in the regulation of the MYC/ERF antagonism upon feeding by *P. rapae*, we determined the effect of exogenously applied ABA on the *P. rapae*-induced expression levels of *VSP2* and *PDF1.2*. Application of 100 µM ABA alone did not significantly activate or repress the expression of *VSP2* or *PDF1.2* in any of the tested lines at any of the tested time points (Figure 4 shows the 30 h time point). Interestingly, caterpillar-induced transcription levels of *VSP2* were significantly enhanced in Col-0, *aba2-1* and *ein2-1* plants at 30 h when ABA was applied to the plants 24 h prior to the start of *P. rapae* infestation (Figure 4). This ABA-mediated enhancement of *P. rapae*-induced *VSP2* expression was not observed in *myc2* and *myc2,3,4* plants. Furthermore, the induction of *MYC2* gene expression by *P. rapae* feeding was blocked in *aba2-1* plants and restored by ABA treatment (Supplemental Figure 1A). This indicates that ABA acts positively on the *P. rapae*-induced MYC-branch, possibly by inducing expression and activity of MYC2. On the other hand, ABA application diminished the high *P. rapae*-induced *PDF1.2* transcript levels in *myc2* and *aba2-1* plants at 30 h. In *myc2,3,4* plants*, PDF1.2* levels were significantly reduced by ABA at 48 h (Supplemental Figure 1B), but not yet significantly at 30 h (Figure 4). This indicates that ABA antagonizes the activation of the ERF-branch independently of the MYC2, MYC3 and MYC4 transcription factors.

**Figure 4:**
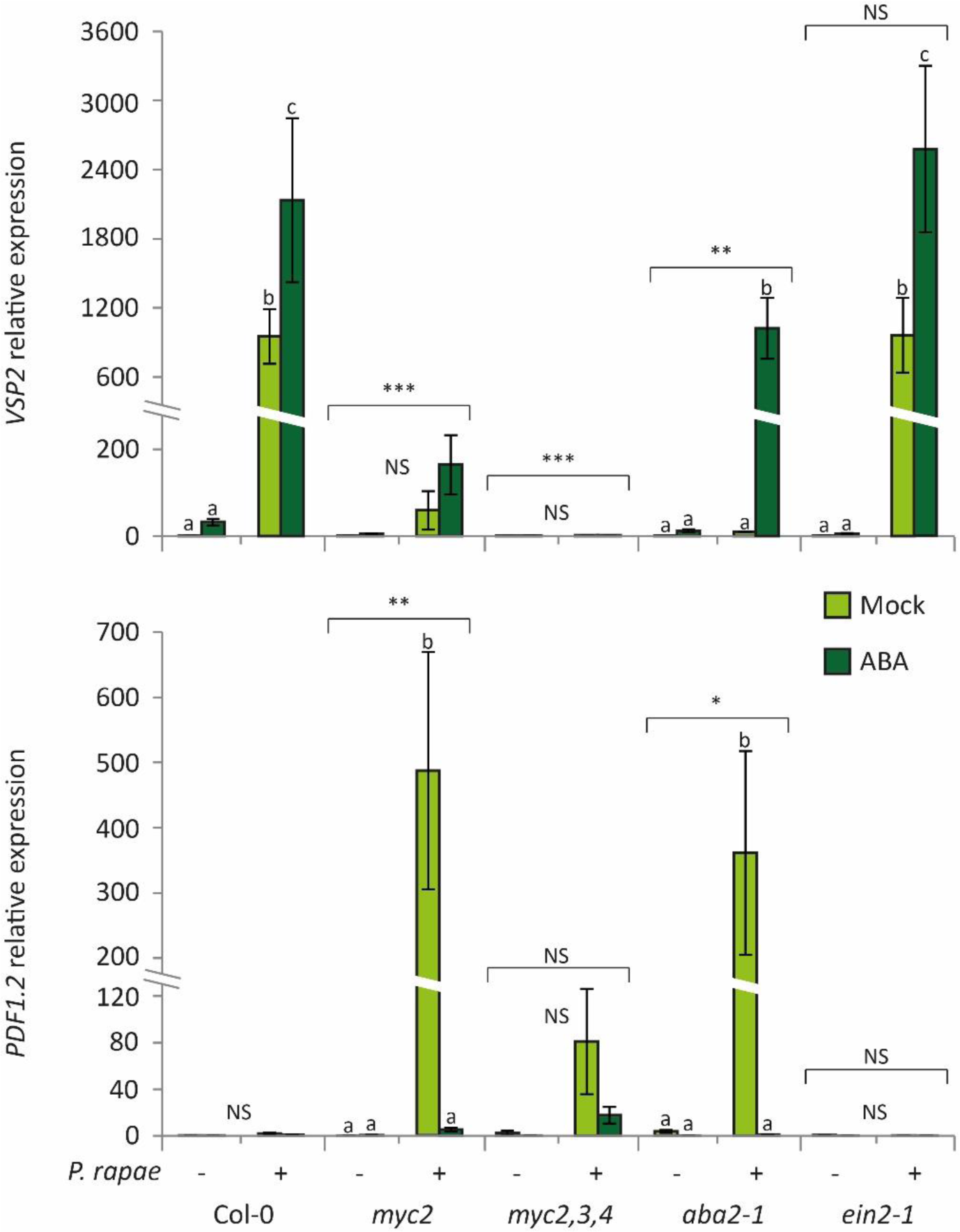
Effect of ABA treatment on *P. rapae*-induced *VSP2* and *PDF1.2* expression. RT-qPCR analysis of *VSP2* and *PDF1.2* gene expression at 30 h in leaves of Col-0, *myc2*, *myc2,3,4*, *aba2-1* and *ein2-1* plants that were treated with a mock solution or with 100 µM ABA 24 h prior to infestation with *P. rapae*. For experimental detail on timing of *P. rapae* treatment, see legend Figure 1. Indicated are expression levels relative to non-infested Col-0 plants at 24 h. Different letters indicate statistically significant differences between treatments of one line. Indications above the brackets specify whether there is an overall statistically significant difference between the mutant line and Col-0 (two-way ANOVA (treatment x genotype), LSD test for multiple comparisons; *** = *P*<0.001; ** = *P*<0.01; * = *P*<0.05; NS = not significant). Error bars represent SE, *n*=3 plants.

ORA59 is a crucial transcription factor for activation of the ERF-branch marker gene *PDF1.2* (Pré et al., 2008). To test if ABA can interfere with *PDF1.2* activation downstream of the ORA59 protein, we used a *35S::ORA59* overexpression line in which *PDF1.2* is constitutively expressed. The *VSP2* expression pattern in the *35S::ORA59* line was similar to that in Col-0 after feeding by *P. rapae* and application of ABA (Figure 5A). *ORA59* levels were constitutively high in the *35S::ORA59* plants and were not significantly influenced by *P. rapae* or ABA treatment. As expected, *PDF1.2* was expressed constitutively in untreated *35S::ORA59* plants and was increased further upon feeding by *P. rapae*, which likely can be ascribed to the elevated JA content in response to herbivory. Application of ABA significantly repressed the *PDF1.2* levels in *P. rapae*-infested *35S::ORA59* plants and a similar trend was found in the non-infested plants (Figure 5A). These results suggest that ABA antagonizes *PDF1.2* expression downstream of ORA59.

**Figure 5:**
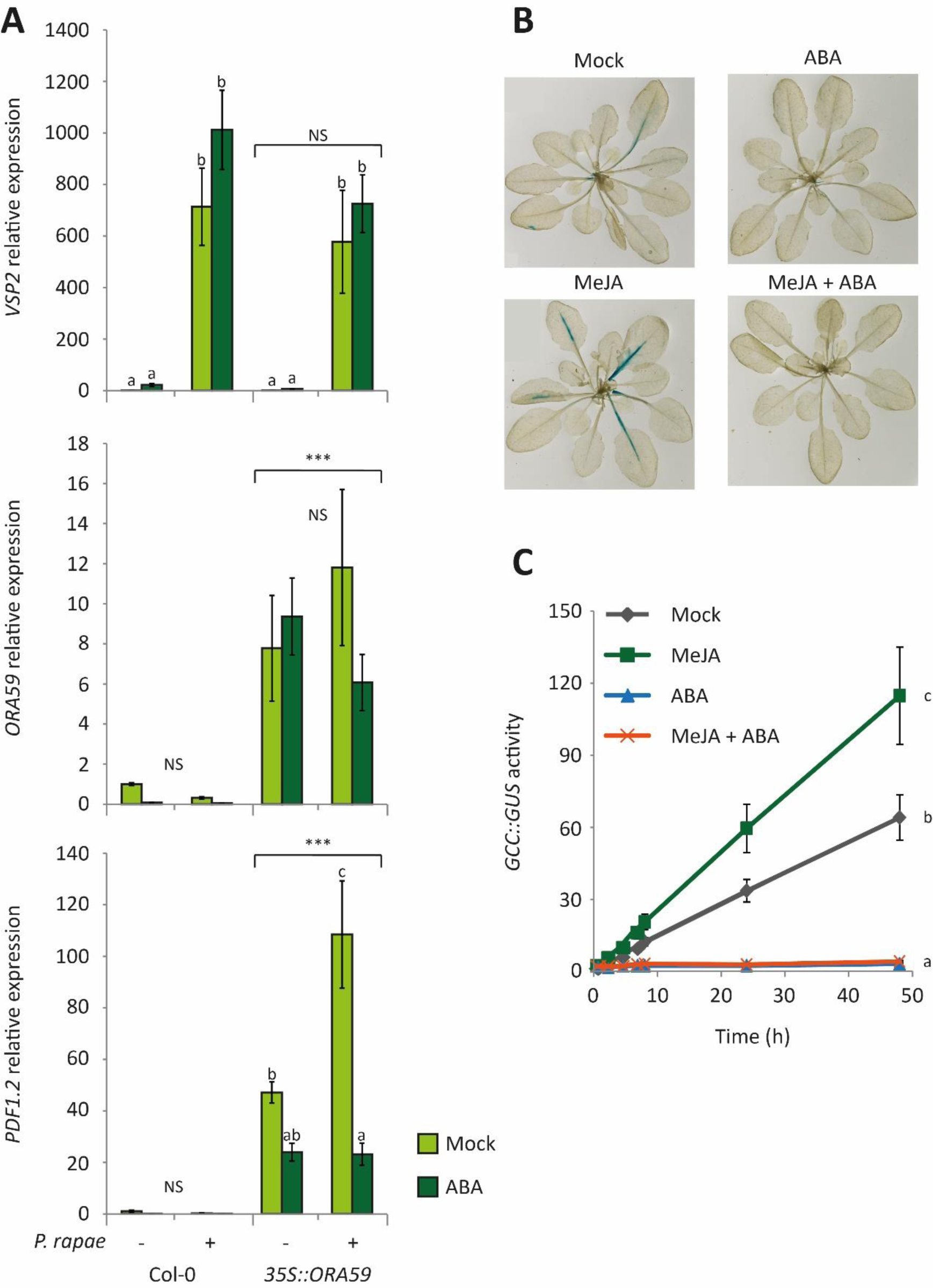
Suppression of *P. rapae*-induced *PDF1.2* expression and MeJA-induced *GCC::GUS* activity by ABA. A) RT-qPCR analysis of *VSP2*, *ORA59* and *PDF1.2* gene expression at 30 h in leaves of Col-0 and *35S::ORA59* plants that were treated with a mock solution or with 100 µM ABA 24 h prior to infestation with *P. rapae*. For experimental detail on timing of *P. rapae* treatment, see legend Figure 1. Indicated are expression levels relative to untreated Col-0 plants at 30 h. Different letters indicate a statistically significant difference between treatments of one line. Indications above the brackets specify whether there is an overall statistically significant difference between *35S::ORA59* and Col-0 (two-way ANOVA (treatment x genotype), LSD test for multiple comparisons; *** = *P*<0.001; NS = not significant). Error bars represent SE, *n*=3 plants. B, C) GUS activity of the *GCC::GUS* line. Plants were dipped in a solution containing 100 µM MeJA, 100 µM ABA, a combination of both chemicals or a mock solution and harvested after 24 h. B) Rosettes were stained for GUS activity or C) GUS activity in the leaves was quantified for 48 h using a microplate reader. Different letters indicate statistically significant differences between treatments (regression analysis; *P*<0.05). Error bars represent SE, *n*=4 plants.

The GCC-box motif that is present in the promoter region of the *PDF1.2* gene and is the binding site for ERF transcription factors has previously been shown to be sufficient for transcriptional activation by JA and suppression thereof by salicylic acid (SA; Brown et al., 2003; Spoel et al., 2003; Van der Does et al., 2013). Therefore, we tested if the GCC-box is also targeted for suppression by ABA. We used a transgenic *GCC::GUS* line containing 4 copies of the GCC-box fused to a minimal *35S* promoter and the *GUS* reporter gene (Zarei et al., 2011). We treated the plants with 100 µM MeJA, 100 µM ABA or a combination of MeJA and ABA and determined GUS activity after 24 h. Both the histochemical staining (Figure 5B) and the quantification of the GUS activity (Figure 5C) showed that MeJA induced GUS activity, confirming that MeJA activates the GCC-box. Treatment with ABA alone significantly repressed the background GUS activity. Moreover, the combination treatment resulted in a significant suppression of the MeJA-induced activation of the GCC-box by ABA. Also the promoter region of the *ORA59* gene contains a GCC-box and in accordance with the *PDF1.2* expression pattern (Figure 4), the high *P. rapae*-induced expression level of *ORA59* in *myc2* plants could be suppressed by prior treatment with ABA (Supplemental Figure 2). Together, these results indicate that, in line with the reported antagonism of JA-dependent gene transcription by SA, the GCC-box is similarly targeted by ABA, which likely contributes to suppression of the ERF-branch of the JA pathway during herbivory.

### The role of ET in regulation of MYC/ERF antagonism

Although the impact of ET signaling on the expression of the MYC- and the ERF-branch upon *P. rapae* feeding is not merely as great as that of ABA, we did observe that in *ein2-1* plants *VSP2* transcription was significantly enhanced at 24 h compared to Col-0 (Figure 1). Furthermore, the production of JA, JA-Ile and ABA was significantly enhanced in *ein2-1* plants at 30 h compared to Col-0 (Figure 2). To investigate whether activation of ET signaling could influence the balance between the MYC- and ERF-branch of the JA pathway during *P. rapae* feeding, we exogenously applied gaseous ET before and during infestation of Col-0 and *ein2-1* plants with *P. rapae* caterpillars. Treatment with 1 ppm of gaseous ET alone induced the expression of *PDF1.2* in Col-0, which was further enhanced by the combination with *P. rapae* feeding (Figure 6). This is likely due to synergistic action between ET and *P. rapae*-induced JAs on *PDF1.2* expression (Penninckx et al., 1998). Additionally, ET treatment strongly reduced the level of both basal and *P. rapae*-induced expression of *VSP1/2*, indicating that induced ET signaling can antagonize the MYC-branch. Both the stimulating effect of ET on *PDF1.2* and the suppressive effect of ET on *VSP1/2* were absent in *P. rapae*-infested *ein2-1* plants, indicating that both ET-mediated processes are dependent on EIN2 and thus regulated via the ET signaling pathway.

**Figure 6:**
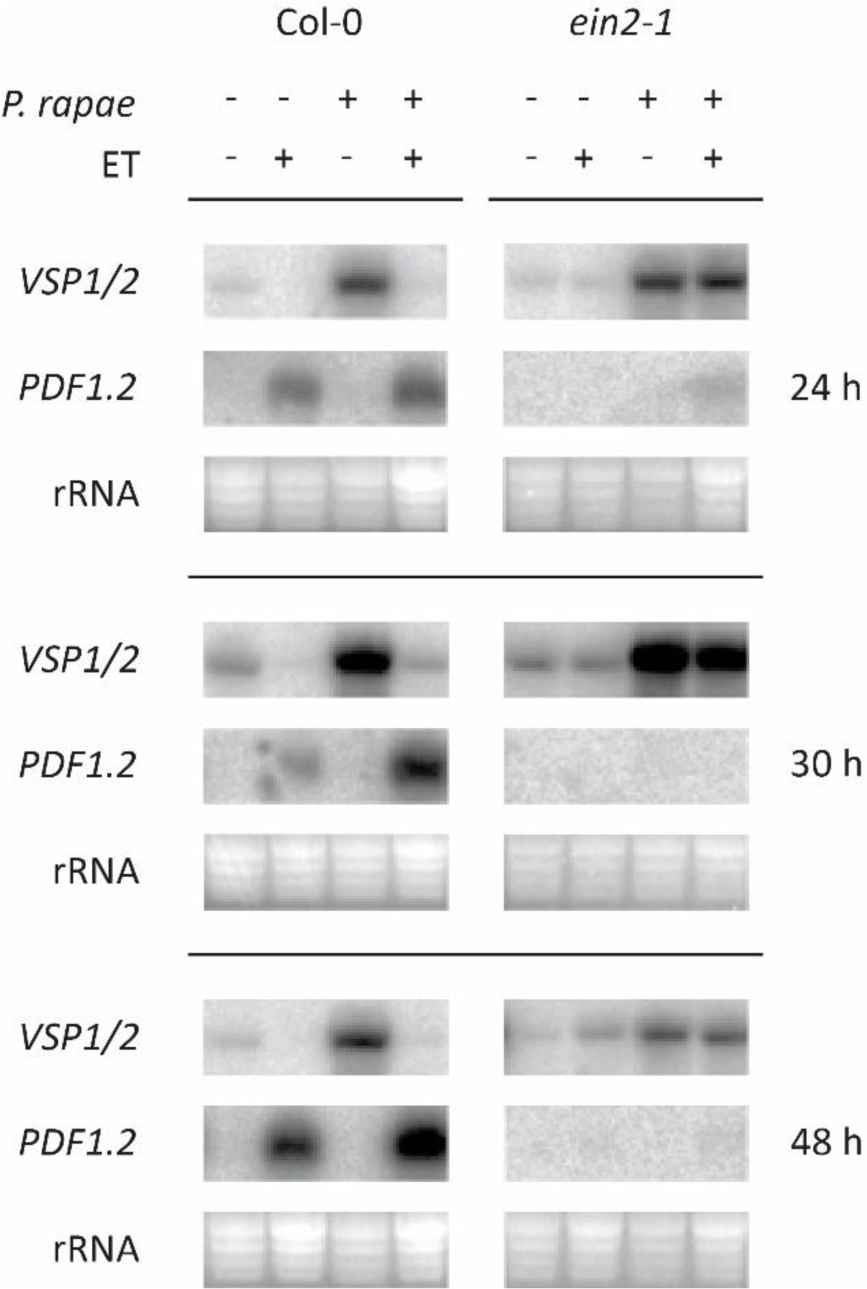
Effect of gaseous ET treatment on *P. rapae*-induced *VSP1/2* and *PDF1.2* expression. Northern blot analysis of *VSP1/2* and *PDF1.2* gene expression in leaves of Col-0 and *ein2-1* plants that were infested with *P. rapae* and treated with a continuous flow of gaseous ET (1 ppm) or ambient air (starting 24 h prior to infestation and continuing until tissue was harvested). First-instar caterpillars of *P. rapae* were allowed to feed for 24 h after which they were removed. Infested leaves were harvested at the indicated time points after *P. rapae* was introduced.

Infection with *B. cinerea* induced ET production (Figure 3B) and we tested if *B. cinerea* infection can also suppress the *P. rapae*-induced activation of the MYC-branch, thereby influencing the MYC/ERF antagonism. Per Col-0 plant, six leaves were inoculated with droplets of *B. cinerea* spores and 24 h later one first-instar caterpillar was placed on the plant. Caterpillars were allowed to feed for 24 h, after which they were removed. *B. cinerea* infection strongly induced the expression of *ORA59* and *PDF1.2* (Figure 7), indicating that the ERF-branch was activated. *P. rapae* infestation activated the MYC-branch as evidenced by enhanced transcription of *VSP2* (Figure 7). Surprisingly, infection with *B. cinerea* prior to *P. rapae* infestation did not antagonize the *P. rapae*-induced activation of *VSP2*. In contrast, *P. rapae* infestation subsequent to *B. cinerea* infection suppressed the *B. cinerea*-induced activation of *ORA59* and *PDF1.2* to basal expression levels (Figure 7). Together, these results suggest that in this set-up Arabidopsis plants prioritize their MYC-branch controlled defenses to combat *P. rapae* infestation, even when the plants were first conditioned to express the ERF-branch defenses against *B. cinerea* infection.

**Figure 7:**
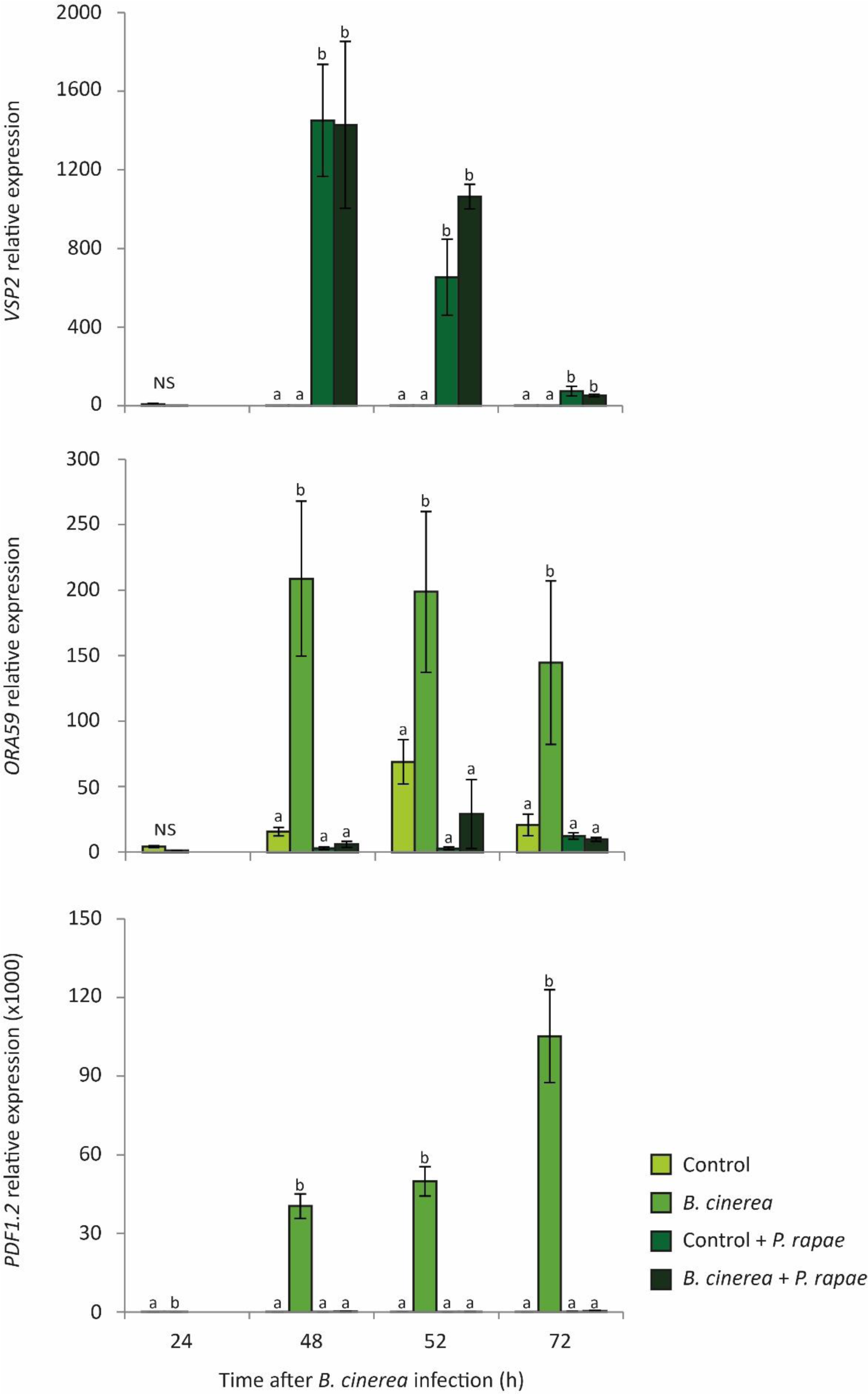
Effect of *B. cinerea* infection on *P. rapae*-induced gene expression. A) RT-qPCR analysis of *VSP2,* ORA59 and *PDF1.2* gene expression in leaves of Col-0 control plants and leaves infected with *B. cinerea* 24 h prior to infestation with *P. rapae*. Indicated are expression levels relative to untreated Col-0 plants at 0 h. Different letters indicate statistically significant differences between the treatments at the indicated time point (ANOVA, Tukey post-hoc tests; *P*<0.05; NS = not significant). Error bars represent SE, *n*=3 plants.

### The effect of ABA and ET on preference and performance of *P. rapae* caterpillars

Previously, Verhage et al. (2011) showed that *P. rapae* caterpillars prefer to feed from plants that express the ERF-branch over plants that express the MYC-branch. Here, we determined the effect of ABA and ET signaling on the preference of *P. rapae* by conducting two-choice assays, in which two plants of each of two genotypes were placed together in a two-choice arena. Leaves were in physical contact with each other, which allowed the caterpillars to freely move from plant to plant. Two first-instar *P. rapae* caterpillars were placed on each plant at the start of the assay (eight caterpillars per arena) and after 4 days the number of caterpillars per plant genotype was determined in 20-30 independent two-choice arenas. As demonstrated previously (Verhage et al., 2011), significantly more caterpillars were detected on *myc2* than on Col-0 plants (Figure 8A). Similarly, *aba2-1* plants contained more caterpillars than Col-0 when tested in a choice assay (Figure 8A). This finding is in accordance with a preference of *P. rapae* caterpillars for plants that express the ERF-branch, as shown for *myc2* and *aba2-1* upon infestation with *P. rapae* (Figure 1). Mutant *ein2-1* plants that, like Col-0, expressed the MYC-branch and not the ERF-branch (Figure 1) accommodated a similar amount of caterpillars as Col-0 plants in a two-choice set-up. These results suggest that MYC2- and ABA-dependent suppression of the ERF-branch in wild-type Col-0 plants during feeding by *P. rapae* reduces the preference of the caterpillars, whereas ET signaling does not influence caterpillar preference.

**Figure 8:**
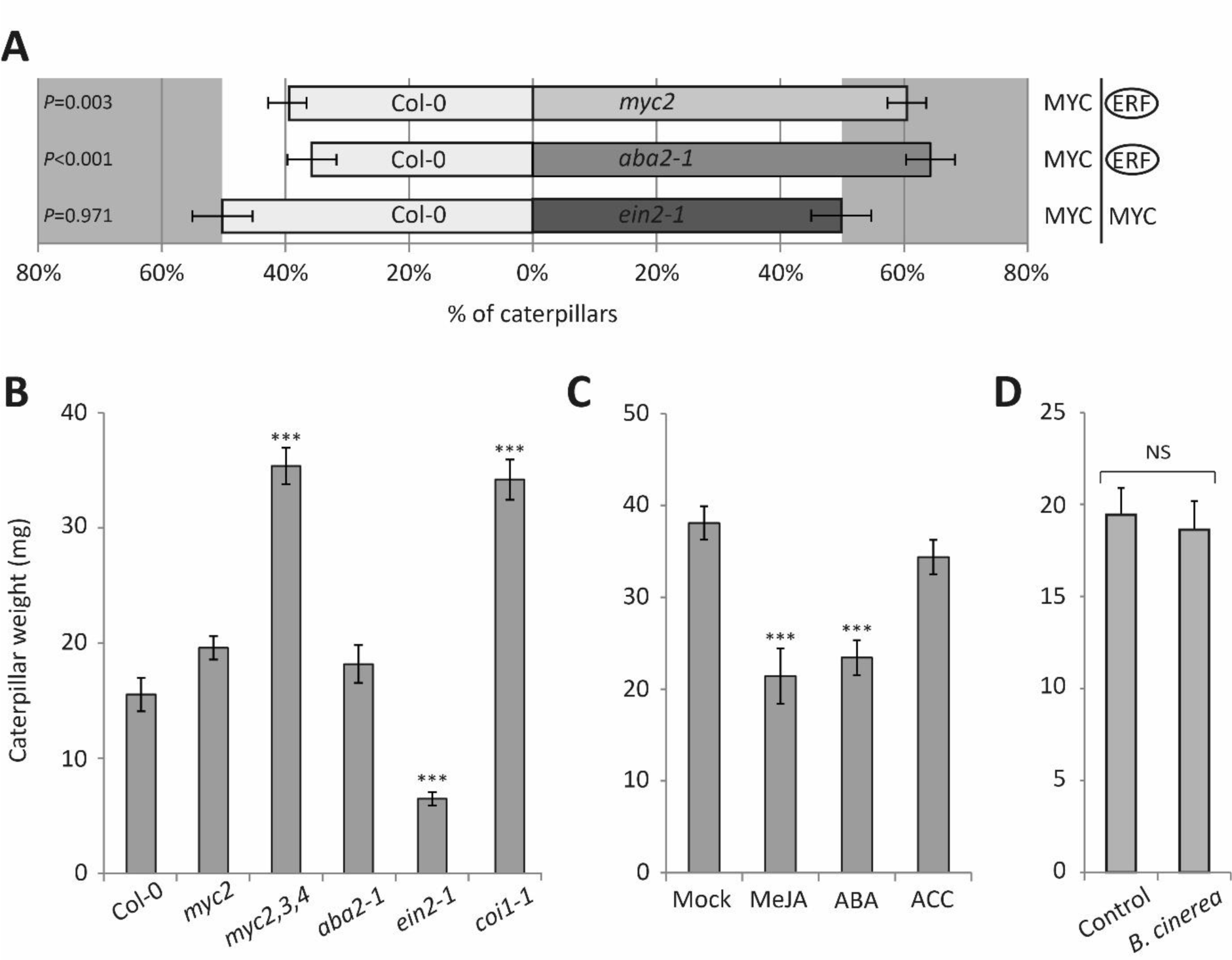
Effect of ABA and ET signaling on the preference and performance of P. rapae. A) Caterpillar preference for Col-0 vs *myc2*, Col-0 vs *aba2-1* and Col-0 vs *ein2-1* plants. Two-choice arenas (*n*=20-30) consisted of two pots per genotype. In each two-choice arena, two first-instar *P. rapae* caterpillars were placed on the plants in each pot (total eight caterpillars per arena). After 4 days the number of caterpillars on each genotype was determined. The right panel displays which branch of the JA pathway is predominantly activated in the corresponding genotypes that are displayed in the left panel. Displayed are the average percentages (±SE) of the distribution of the *P. rapae* caterpillars over the two genotypes (x-axis). *P-*values indicate a statistically significant difference from the 50% percentile (Student’s *t*-test). In cases of statistically significant differences (*P*<0.05), the preferred branch of the JA pathway is marked with a circle. Experiments were repeated with similar results. B, C, D) Caterpillar performance on Col-0, *myc2*, *myc2,3,4*, *aba2-1*, *ein2-1* and *coi1-1* plants (B), on Col-0 plants treated with a mock solution, 100 µM MeJA, 100 µM ABA or 1 µM ACC (C) and on control plants and plants treated with *B. cinerea* (D). The hormone solutions were applied as root-drench at 5 and 2 days before caterpillar feeding. One first-instar *P. rapae* caterpillar was placed on each plant and allowed to feed for 7 days after which the weight was determined. Asterisks indicate a statistically significant difference in comparison to Col-0 or mock-treated plants (ANOVA, Dunnett post-hoc tests; *** = *P*<0.001; * = *P*<0.05, NS = not significant). Error bars represent SE, *n*=8-28 plants.

To investigate whether the preference of *P. rapae* caterpillars for the ERF-branch-expressing *myc2* and *aba2-1* mutant plants coincides with increased performance of the caterpillars on these genotypes, we assessed their growth in no-choice assays with Col-0, *myc2*, *myc2,3,4*, *aba2-1*, *ein2-1*, and JA-nonresponsive *coi1-1* plants. One first-instar *P. rapae* caterpillar was placed on each plant and allowed to feed for 7 days, after which the caterpillar was weighed. Figure 8B shows that there was no significant difference between the growth of caterpillars that fed from Col-0, *myc2* or *aba2-1*. In contrast, on *ein2-1* mutants, caterpillar growth was significantly inhibited. The growth of caterpillars on *myc2,3,4* mutants was increased to the same extent as on *coi1-1* mutants. Next, we tested whether pretreatment of Col-0 plants with solutions of 100 µM MeJA, 100 µM ABA or 1 µM of the ET precursor 1-aminocyclopropane-1-carboxylic acid (ACC) had an effect on caterpillar performance. MeJA or ABA pretreatment significantly reduced the weight of the caterpillars, whereas pretreatment with ACC did not have an effect (Figure 8C). Finally, the effect of prior infection with ET-inducing *B. cinerea* on *P. rapae* performance was tested. Figure 8D shows that *P. rapae* performance was not altered on *B. cinerea*-infected plants compared to control plants.

These results indicate that, although caterpillars have a preference for the ERF-branch-expressing *myc2* and *aba2-1* plants, there is no direct positive effect on their performance by these plants. In accordance, the ERF-branch-activating ACC and *B. cinerea* pretreatments had no effect on caterpillar performance. On the other hand, enhanced activation of the MYC-branch as is evident in *ein2-1* plants upon caterpillar feeding (Figure 1) correlates with reduced performance of the caterpillars on these plants. Moreover, also the MYC-branch-activating/priming MeJA and ABA pretreatments significantly reduced caterpillar performance (Figure 8C). In conclusion, enhancement of the MYC-branch, by activating the JA or ABA pathway or by suppressing the ET pathway reduced caterpillar performance. The preference for plants expressing the ERF-branch, as is demonstrated in the two-choice assays, might be a strategy to avoid plants with an effective defense.

## Discussion

The complex plant immune regulatory network that is activated upon recognition of attackers is largely controlled by plant hormones (Pieterse et al., 2012). JA has a decisive regulatory role in the defense responses against herbivorous insects and necrotrophic pathogens (Howe and Jander, 2008; Pieterse et al., 2012). Several studies indicated that ABA co-regulates the JA-induced activation of the MYC-branch, while ET co-regulates activation of the ERF-branch (Penninckx et al., 1998; Lorenzo et al., 2003; Anderson et al., 2004; Lorenzo et al., 2004; Pré et al., 2008; Vos et al., 2013b). Previously, Verhage et al. (2011) showed that feeding of *P. rapae* caterpillars on Arabidopsis leads to activation of the MYC-branch while the herbivore-preferred ERF-branch is strongly suppressed. However, the role of ABA and ET in the antagonistic interaction between the MYC- and the ERF-branch during herbivory was not clear. Here, we show that ABA and ET are important regulators of the balance between the MYC- and the ERF-branch in herbivore-infested plants, thereby activating the appropriate defense response and suppressing costly unnecessary defenses.

### ABA is required for *P. rapae*-induced activation of the MYC-branch and repression of the ERF-branch

There is ample evidence for the production of JA upon feeding by chewing herbivores (Wasternack and Hause, 2013), but the production of ABA is not often taken along. We demonstrated that in wild-type Col-0 plants, *P. rapae* feeding enhanced the production of JAs, as well as that of ABA (Figure 2 & Figure 9; Vos et al., 2013b). Also in maize and rice plants, an increased production of JAs and ABA has been demonstrated upon root herbivory (Erb et al., 2009; Lu et al., 2015). We show that *aba2-1* plants fail to activate the MYC-branch in response to *P. rapae* feeding, evidenced by the reduced activation of *MYC2* and the MYC-branch marker *VSP2* (Figure 1 and Supplemental Figure 1A). Importantly, *aba2-1* plants are also deficient in suppression of the ERF-branch in response to herbivory, apparent from enhanced activation of *PDF1.2* after *P. rapae* feeding (Figure 1). Since *aba2-1* plants differ from Col-0 in the herbivory-induced production of ABA and only minimally in the production of JAs (Figure 2), it seems plausible that in wild-type Arabidopsis ABA is essential for shifting the MYC/ERF balance towards the MYC-branch upon herbivory by *P. rapae*. This was confirmed by experiments in which ABA was applied exogenously 24 h prior to infestation with *P. rapae*. The ABA treatment stimulated the herbivore-induced MYC-branch in Col-0 plants, while in *myc2* and *myc2,3,4* plants ABA treatment strongly inhibited the enhanced expression of the ERF-branch (Figure 4 & Supplemental Figure 1B). In line with this, induction of the ERF-branch by *B. cinerea* infection was strongly suppressed by subsequent *P. rapae* feeding (Figure 7), likely due to enhanced ABA levels in response to *P. rapae* feeding (Vos et al., 2015). Treatment with exogenous ABA in the absence of herbivory did not alter the expression of the marker genes *VSP2* and *PDF1.2* (Figure 4), indicating that ABA alone is not sufficient for influencing the expression levels of these marker genes, but requires additional activation of the JA pathway. Likewise, we previously demonstrated that systemic induction of *MYC2* in non-damaged leaves of *P. rapae*-infested plants only led to downstream activation of *VSP2* if ABA levels increased as well (Vos et al., 2013b).

**Figure 9:**
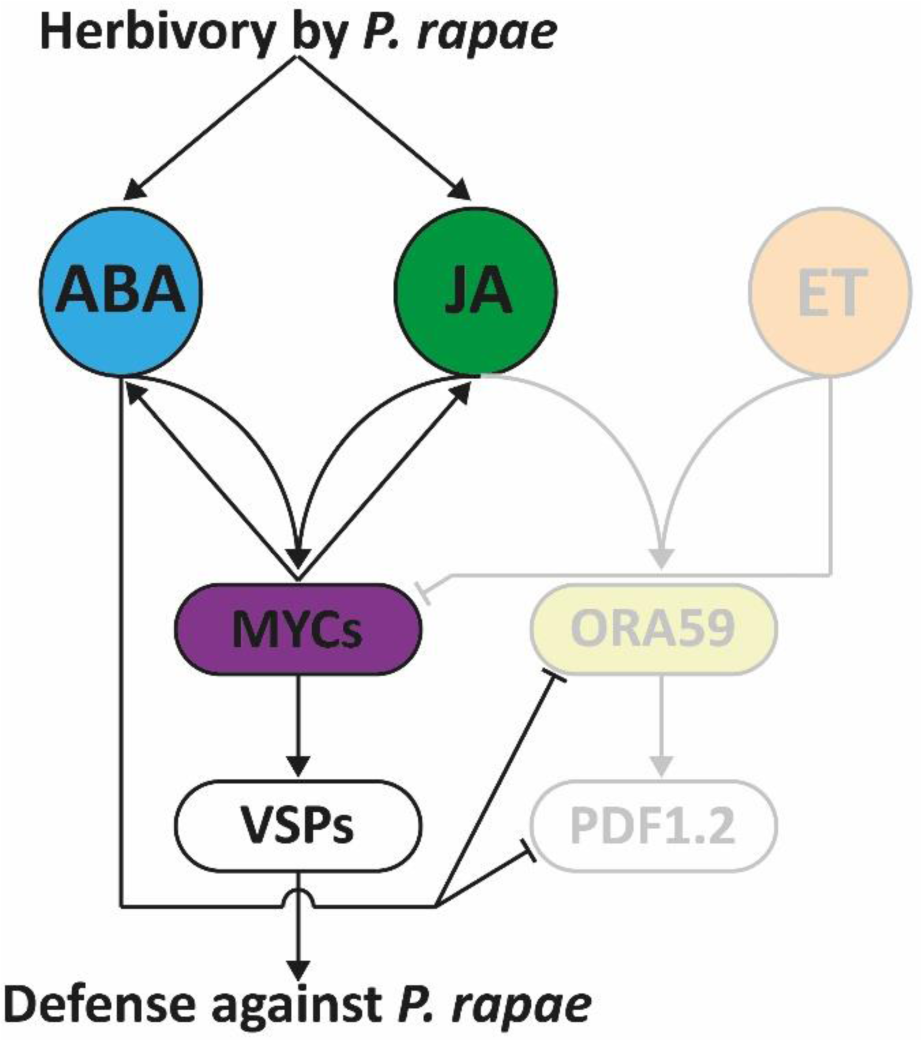
Model of differential regulation of JA responses during herbivory by P. rapae. Feeding by *P. rapae* stimulates the production of JAs and ABA, resulting in activation of the MYC-branch and a concomitant defense response against *P. rapae*. While the activation of the MYC transcription factors is dependent on ABA, the production of ABA is dependent on the MYC transcription factors, resulting in a positive feedback loop between ABA and the MYC2, MYC3 and MYC4 transcription factors. Simultaneously, activation of the ABA pathway suppresses the ERF-branch at the level of transcriptional activation at the GCC-box. The ERF-branch components indicated in the model are shaded, because they are not activated during the Arabidopsis-*P. rapae* interaction. Although ET has the capacity to suppress the *P. rapae*-induced MYC-branch, it is not produced during the Arabidopsis-*P. rapae* interaction, and thus does not play a significant role in the MYC/ERF-branch interaction model during infestation. Arrows indicate a stimulating effect, blocked lines indicate a suppression.

Interestingly, the suppression of the ERF-branch by exogenous application of ABA to *P. rapae*-infested plants occurred in the *myc2* and *myc2,3,4* plants (Figure 4 & Supplemental Figure 1), showing that this response is independent of the previously reported MYC2/MYC3/MYC4-EIN3/EIL1 protein-protein interactions (Song et al., 2014a). However, the ABA biosynthesis that was induced upon *P. rapae* feeding was largely dependent on MYC transcription factors, as indicated by basal ABA levels in *P. rapae-*infested *myc2* and *myc2,3,4* plants compared with Col-0 at 24 h (Figure 2). Also in Arabidopsis roots JA-dependent signaling was reported to be necessary for the production of ABA (De Ollas et al., 2015). Hence, despite the fact that exogenous ABA can suppress ERF-branch-induced responses independently of the MYC transcription factors, there seems to be a positive feedback loop between the ABA and JA response pathways that are induced by herbivory, in which ABA signaling enhances MYC transcription factor activity, which in turn is important for the production of ABA (Figure 9).

### ABA antagonizes the ERF-branch downstream of ORA59 at the GCC-box

Analysis of the *35S::ORA59* transgenic line showed that ABA is able to suppress *PDF1.2* even when ectopic *ORA59* expression levels are constitutively high (Figure 5A). Previously, Van der Does et al. (2013) investigated the suppressive effect of SA on JA-induced *PDF1.2* expression. They also found that SA could suppress activation of *PDF1.2* in the *35S::ORA59* line. Moreover, they reported that the GCC-box, which is present in the promoter of *PDF1.2*, and required for the JA-responsive expression, is essential and sufficient for transcriptional suppression by SA. Similarly, we show here that ABA strongly inhibits activation of the GCC-box in mock- or MeJA-treated plants (Figure 5B & C). In addition to *PDF1.2* also *ORA59* harbors a GCC-box in its promoter, and activated expression of *ORA59* is shown to be suppressed by (*P. rapae*-induced) ABA (Figure 4, Figure 7 & Supplemental Figure 2), as well as by SA (Zander et al.,2014). Together, these data point towards a similar mechanism for SA-dependent and ABA-dependent suppression of the expression levels of *ORA59* and *PDF1.2* at the level of transcriptional regulation at the GCC-box.

### Strong activation of the ET pathway is necessary for suppression of the MYC-branch

Continuous monitoring of the production of ET in *P. rapae-*infested Arabidopsis plants revealed no changes in the emission of ET in this set-up (Figure 3 & Figure 9). However, ET-nonresponsive *ein2-1* plants showed enhanced activation of the MYC-branch, as evidenced by an increase in *VSP2* transcription at 24 h after introduction of *P. rapae* (Figure 1). The production of JA-Ile, JA and especially ABA was enhanced in the *ein2-1* plants compared with Col-0 upon *P. rapae* feeding (Figure 2), suggesting that in wild-type plants basal activity of the ET pathway can inhibit herbivory-induced production of JA and ABA, which tempers the activation of the MYC-branch.

Strong evidence for a role for ET in shifting the MYC/ERF balance was provided by the experiment in which we applied gaseous ET to the plants. This ET treatment led to activation of the ERF-branch during *P. rapae* feeding, while the MYC-branch was suppressed (Figure 6). Both effects were absent in the ET-insensitive mutant *ein2-1,* indicating that the modulating effect of ET was mediated via the ET pathway. Infection with the necrotrophic pathogen *B. cinerea* induced ET emission (Figure 3), but in contrast to exogenously applied gaseous ET, *B. cinerea* infection was not able to suppress activation of the MYC-branch in response to *P. rapae* feeding (Figure 7). In a previous study, where *P. rapae* feeding preceded *B. cinerea* infection, the MYC-branch was not suppressed either (Vos et al., 2015). A likely explanation for the discrepancy between the continuous application of gaseous ET and infection with *B. cinerea* is that only the gaseous ET treatment activates the ET pathway to a great enough extent to suppress the *P. rapae*-induced MYC-branch.

ET has been described to play a role in resistance to herbivores in many plant species (Von Dahl and Baldwin, 2007), for example as a volatile signal (Broekgaarden et al., 2015). However, the specific effect of ET on plant defense varies per plant species and attacking insect (Nguyen et al., 2016). Although ET has the potential to modulate the balance between the MYC- and the ERF-branch in Arabidopsis, ET levels do not change upon feeding by *P. rapae* only excessive amounts of ET seem to be able to suppress the MYC-branch upon *P. rapae* feeding. Therefore, the induction of the ET pathway is unlikely to play a major role in the defense response of Arabidopsis to *P. rapae* feeding (Figure 9).

### The differential role of the MYC- and the ERF-branch on preference and performance of *P. rapae* caterpillars

The importance of ABA in defense against insect herbivory became further apparent from bioassays in which preference and performance of *P. rapae* caterpillars was determined. In two-choice assays *P. rapae* caterpillars were found to prefer to feed from the *aba2-1* and *myc2* plants over wild-type Col-0 plants (Figure 8A). These findings confirm previous results that *P. rapae* caterpillars prefer to feed from plants expressing the ERF-branch (Verhage et al., 2011). There was no preference for either *ein2-1* or Col-0 plants when caterpillars were given a choice between those two genotypes in a two-choice assay (Figure 8A). Comparably, the specialist caterpillar *P. xylostella* did not move away faster from leaves in which the MYC-branch was activated compared to control leaves (Perkins et al., 2013). The feeding preference of *P. rapae* caterpillars for *aba2-1* and *myc2* plants was not obviously correlated with enhanced performance (weight gain) on these mutants in no-choice assays (Figure 8B), which corresponds with the observation that the ERF-branch activating *B. cinerea* infection or ACC pretreatment did not affect caterpillar performance (Figure 5C & 5D; Davila Olivas et al.,2016). Performance of both specialist *P. rapae* caterpillars (Figure 8B) and generalist *S. littoralis* caterpillars (Bodenhausen and Reymond, 2007) was highly reduced on *ein2-1* plants, which corresponds with the observation that the MYC-branch-activating/priming MeJA or ABA treatment significantly reduced caterpillar performance (Figure 8C). Moreover, caterpillar performance on the *myc2,3,4* and *coi1-1* mutants was greatly enhanced (Figure 8B). This indicates that full activation of the MYC-branch is needed to effectively reduce growth of the caterpillars, while activation of the ERF-branch determines preference of specialist caterpillars, which may enable them to choose plants on which their performance is not negatively affected.

Altogether, this study highlights the interplay between JA on the one hand, and ABA and ET on the other hand, in shaping the outcome of the defense response that is triggered upon caterpillar feeding. We show that induced ABA can activate the MYC-branch upon *P. rapae* feeding and suppress the ERF-branch by targeting the GCC-box. Although ET is also capable of suppressing the MYC-branch, ET production is not induced by *P. rapae* feeding and is not likely to play a role in the defense response upon feeding. By prioritizing the MYC-branch over the ERF-branch during insect herbivory, Arabidopsis is capable of prioritizing its JA-induced response to defenses that contribute to maximizing the chance of survival.

## Material and Methods

### Plant material and cultivation

Seeds of *Arabidopsis thaliana* accession Col-0 and mutants *jin1-7* (*myc2*), *myc2,3,4*, *aba2-1*, *ein2-1* and *coi1-1* (Koornneef et al., 1982; Feys et al., 1994; Alonso et al., 1999; Lorenzo et al., 2004; Fernández-Calvo et al., 2011) and the transgenic lines *35S::ORA59* and *GCC::GUS* (Pré et al., 2008; Zarei et al., 2011) were sown on river sand. Two weeks later, seedlings were transplanted into 60-ml pots containing a sand-potting soil mixture (5:12 v/v) that had been autoclaved twice for 20 min with a 24 h interval. Plants were cultivated in a growth chamber with a 10-h day and 14-h night cycle at 70% relative humidity and 21°C. Plants were watered every other day and received 10 ml of half-strength Hoagland solution (Hoagland and Arnon, 1938) containing 10 µM sequestreen (CIBA-Geigy, Basel, Switzerland) once a week.

### Pieris rapae assays

*Pieris rapae* (small cabbage white) was reared on white cabbage plants (*Brassica oleracea*) as described (Van Wees et al., 2013). First-instar caterpillars were used in all experiments. For gene expression analysis, two caterpillars were placed on fully expanded leaves of 5-week-old Arabidopsis plants using a fine paintbrush. Caterpillars were removed 24 h later and leaves were harvested at different time points after infestation. For the ET production measurement, caterpillars remained on the leaves for the entire assay.

For the two-choice assays, two or three *aba2-1* and *ein2-1* mutant plants (instead of one plant), were grown in one pot to compensate for their smaller size. Biomass and leaf area were measured from a representable subset of 6-week-old plants before the start of the assay to verify that the amount of leaf tissue was equal among the different genotypes tested. Two pots containing Col-0 wild-type plants and two pots a mutant were placed together in an arena, such that there was physical contact between the plants. Two first-instar caterpillars were released on the plants in each pot in the arena (n=20-30), so that there were eight caterpillars per arena that could freely move through the arena. After 4 days, the number of caterpillars present on each genotype was monitored and the frequency distribution of the caterpillars over the different genotypes was calculated.

To examine caterpillar performance, a single first-instar caterpillar was placed on a 5-week-old plant inside a plastic cup covered with an insect-proof mesh to contain the caterpillars. After 7 days of feeding, caterpillars were weighed to the nearest 0.1 mg on a microbalance.

### Botrytis cinerea inoculation

*Botrytis cinerea* inoculations were performed with strain B05.10 (Van Kan et al., 1997) as described previously (Van Wees et al., 2013). *B. cinerea* solution was made into a final density of 1·10^5^ spores/ml and 5 µL droplets of the spores were applied to six leaves of 5-week-old plants. Plants were used immediately for measurement of ethylene production or were placed under a lid for 24 h to increase relative humidity and stimulate the infection, after which the lids were removed and *P. rapae* caterpillars were placed on the plants.

### Chemical treatments

For gene expression analysis, plants were treated with MeJA (Serva, Brunschwig Chemie, Amsterdam, the Netherlands) or ABA (Sigma, Steinheim, Germany) by dipping the rosettes in a solution containing either 100 µM MeJA, 100 µM ABA or a combination of both chemicals and 0.015% (v/v) Silwet L77 (Van Meeuwen Chemicals BV, Weesp, the Netherlands) 24 h before caterpillar feeding. MeJA and ABA solutions were diluted from a 1000-fold concentrated stock in 96% ethanol. The mock solution contained 0.015% Silwet L77 and 0.1% ethanol.

For analysis of caterpillar performance, plants were treated with 100 µM MeJA, 100 µM ABA or 1 µM ET precursor 1-aminocyclopropane-1-carboxylic acid (ACC; Sigma, Steinheim, Germany) by applying 20 ml of the solutions to the plants as a root drench, 5 and 2 days before introduction of the *P. rapae* caterpillars. MeJA and ABA solutions were diluted from a 1000-fold concentrated stock in 96% ethanol. The mock solution contained 0.1% ethanol.

Treatment with gaseous ET was performed as described previously (Millenaar et al., 2005). In short, gaseous ET (100 μl/l; Hoek Loos, Amsterdam, the Netherlands) and air (70% relative humidity) were mixed using flow meters (Brooks Instruments, Veenendaal, the Netherlands) to generate an output concentration of 1 μl/l ET, which was flushed continuously through glass cuvettes (13.5 x 16.0 x 29.0 cm) at a flow rate of 75 l/h and then vented to the outside of the building. The concentration of ET in the airflow was verified using gas chromatography. Five-week-old plants were placed separately in the cuvettes and remained there for the duration of the experiment. Control plants were placed in cuvettes which were flushed with air (70% relative humidity) at the same flow rate. ET and air treatments started 1 day prior to introduction of *P. rapae* to the plants in the cuvettes and continued for the duration of the experiment. Light and temperature conditions were the same as described above.

### RNA extraction, RT-qPCR and northern blot analysis

Total RNA was isolated as described (Oñate-Sánchez and Vicente-Carbajosa, 2008). RevertAid H minus Reverse Transcriptase (Fermentas) was used to convert DNA-free total RNA into cDNA. PCR reactions were performed in optical 384-well plates (Applied Biosystems) with an ABI PRISM® 7900 HT sequence detection system using SYBR ® Green to monitor the synthesis of double-stranded DNA. A standard thermal profile was used: 50°C for 2 min, 95°C for 10 min, 40 cycles of 95°C for 15 s and 60°C for 1 min. Amplicon dissociation curves were recorded after cycle 40 by heating from 60 to 95°C with a ramp speed of 1.0°C/min. Transcript levels were calculated relative to the reference gene At1g13320 (Czechowski et al., 2005) using the 2^-ΔΔCT^ method described previously (Livak and Schmittgen, 2001; Schmittgen and Livak, 2008).

For northern blot analysis, 15 µg of RNA was denatured using glyoxal and dimethyl sulfoxide (Sambrook et al., 1989), electrophoretically separated on 1.5% agarose gel, and blotted onto Hybond-N^+^ membranes (Amersham, ‘s-Hertogenbosch, the Netherlands) by capillary transfer. The electrophoresis and blotting buffer consisted of 10 and 25 mM sodium phosphate (pH 7.0), respectively. Equal loading was confirmed by staining rRNA bands with ethidium bromide. Northern blots were hybridized with gene-specific probes for *PDF1.2* and *VSP1/2* (Leon-Reyes et al., 2010). After hybridization with α-^32^P-dCTP-labeled probes, blots were exposed for autoradiography.

The AGI numbers of the studied genes are At1g32640 (*MYC2*), At5g24780 (*VSP1*), At5g24770 (*VSP2*), At1g06160 (*ORA59*) and At5g44420 (*PDF1.2*).

### Jasmonates and ABA analysis

For JA, JA-Ile, OPDA and ABA concentration analysis, 50-100 mg of *P. rapae*-infested damaged leaves as well as undamaged leaves from non-infested control plants were grinded. The extraction and hormone analysis was performed as previously described (López-Ráez et al., 2010). At the start of the extraction 1 ml of cold ethylacetate containing D6-SA (25 ng/ml) and D5-JA (25 ng/ml) was added to the samples as an internal standard in order to calculate the recovery of the hormones measured. Hormone levels were analyzed by LC-MS on a Varian 320 Triple Quad LC/MS/MS. Ten µl of each sample was injected onto a Pursuit column (C18; 5 µm, 50 x 2.0 mm; Varian) that was connected to a precolumn (Pursuit Metaguard C18; 5 µm; 2.0 mm). Multiple reaction monitoring was performed for parent-ions and selected daughter-ions after negative ionization: JA 209/59 (fragmented under 12V collision energy), JA-Ile 322/130 (fragmented under 19V collision energy), OPDA 291/165 (fragmented under 18V collision energy) and ABA 263/153 (fragmented under 9V collision energy). The mobile phase comprised solvent A (0.05% formic acid) and solvent B (0.05% formic acid in MeOH) with settings as described (Diezel *et al*., 2009). The retention time of each compound was confirmed with pure compounds (ChemIm Ltd, Olomouc, Czech Republic). The surface area for each daughter-ion peak was recorded for the detected analytes. Analytes were quantified using standard curves made for each individual compound.

### Ethylene measurements

ET production was measured in a laser-driven photoacoustic detection system (ETD-300, Sensor Sense, Nijmegen, the Netherlands) connected to a 6-channel valve control box in line with a flow-through system (Voesenek et al., 1990). Five-week-old plants were placed in 2-l air-tight cuvettes (four plants per cuvette), which were incubated under growth chamber conditions. After an acclimation time of 2 h, the cuvettes were continuously flushed with air (flow rate: 0.9 l/h), directing the flow-through air from the cuvettes into a photoacoustic cell for ET measurements. ET levels were measured over consecutive 0.5 h time intervals, after which the machine switched to the next cuvette (n=6).

### GUS assays

For the histochemical GUS assay, GUS activity was assessed by transferring plants to a GUS staining solution (1 mM X-Gluc, 100 mM NaPi buffer, pH 7.0, 10 mM EDTA and 0.1% [v/v] Triton X-100). After vacuum infiltration and overnight incubation at 37°C, the plants were destained by repeated washes in 96% ethanol (Spoel et al., 2003). For the quantitative GUS assay, protein was isolated from frozen plant material and GUS activity was quantified using a microplate reader (BioTek Instruments, Inc., Winooski, United States of America) as described (Pré et al., 2008).

## Acknowledgements

The authors thank Hans Van Pelt and Silvia Coolen for rearing of *P. rapae*, Rob Welschen and Ronald Pierik for their help with experiments on ET application and measurement, Michel de Vries for running the LC/MS/MS samples, Roberto Solano for kindly providing the *myc2,3,4* seeds and Colette Broekgaarden for valuable comments on the manuscript. This research was supported by VIDI grant no. 11281 of the Dutch Technology Foundation STW (to S.C.M.V.W.), VICI grant no. 865.04.002 of the Earth and Life Sciences Foundation, which are part of the Netherlands Organization of Scientific Research (NWO), and ERC Advanced Investigator Grant no. 269072 of the European Research Council (to C.M.J.P.).

## Author contributions

I.A.V., A.V., C.M.J.P. and S.C.M.V.W. designed the research. I.A.V., A.V., L.G.W. and I.V. performed the research. I.A.V., A.V., L.G.W., I.V., R.C.S., C.M.J.P. and S.C.M.W. analyzed the data. I.A.V., C.M.J.P. and S.C.M.W. wrote the paper.

---

**Supplemental Figure 1:**
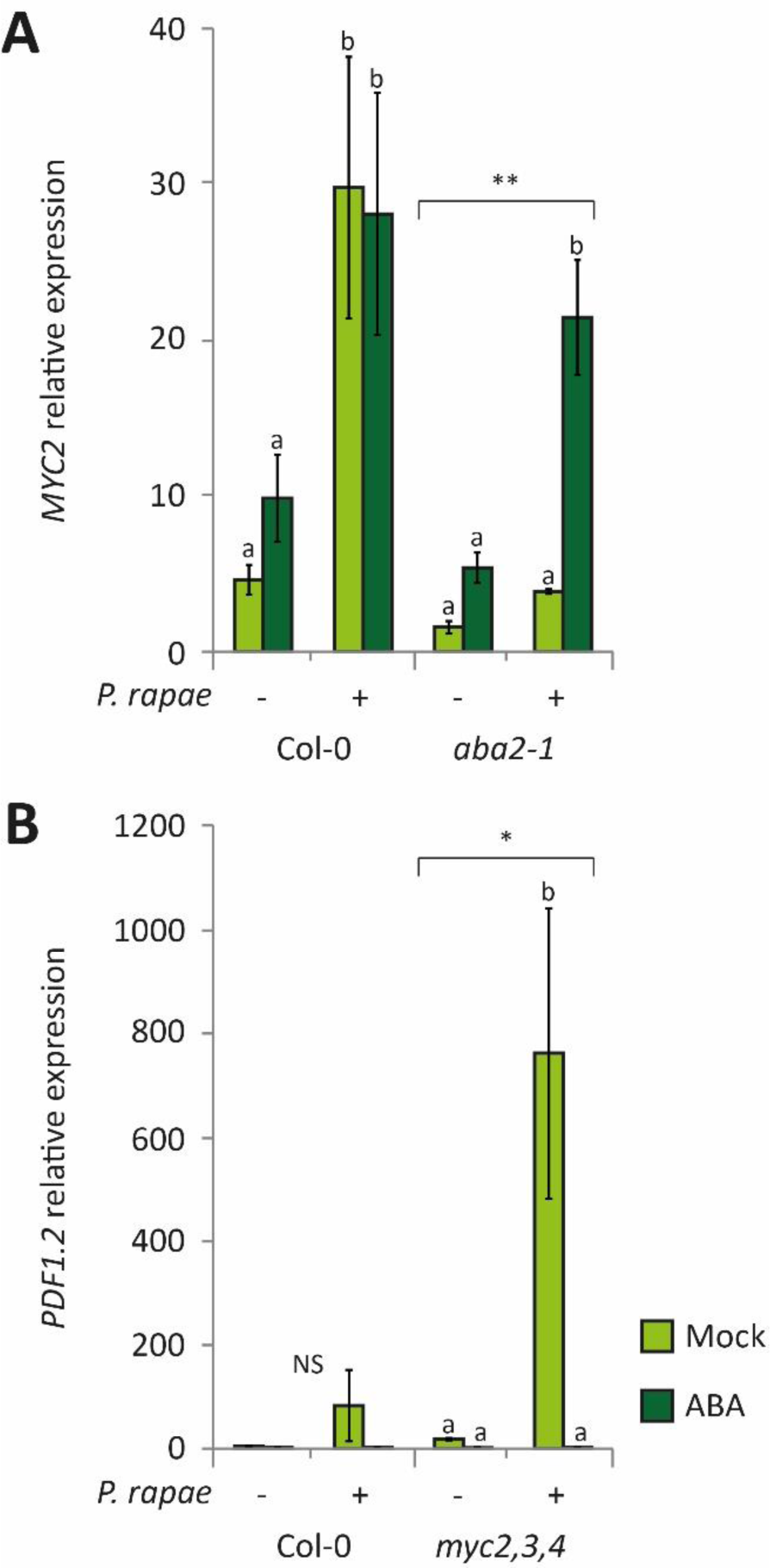
Effect of ABA treatment on *P. rapae*-induced *MYC2* and *PDF1.2* expression. RT-qPCR analysis of *MYC2* expression at 30 h in Col-0 and *aba2-1* plants (A) and *PDF1.2* gene expression at 48 h in leaves of Col-0 and *myc2,3,4* plants (B) that were treated with a mock solution or with 100 µM ABA 24 h prior to infestation with *P. rapae*. For experimental detail on timing of *P. rapae* treatment, see legend Figure 1. Indicated are expression levels relative to non-infested Col-0 plants at 24 h. Different letters indicate statistically significant differences between treatments of one line. Indications above the brackets specify whether there is an overall statistically significant difference between *aba2-1*/*myc2,3,4* and Col-0 (two-way ANOVA (treatment x genotype), LSD test for multiple comparisons; ** = *P*<0.01; * = *P*<0.05; NS = not significant). Error bars represent SE, *n*=3 plants.

**Supplemental Figure 2:**
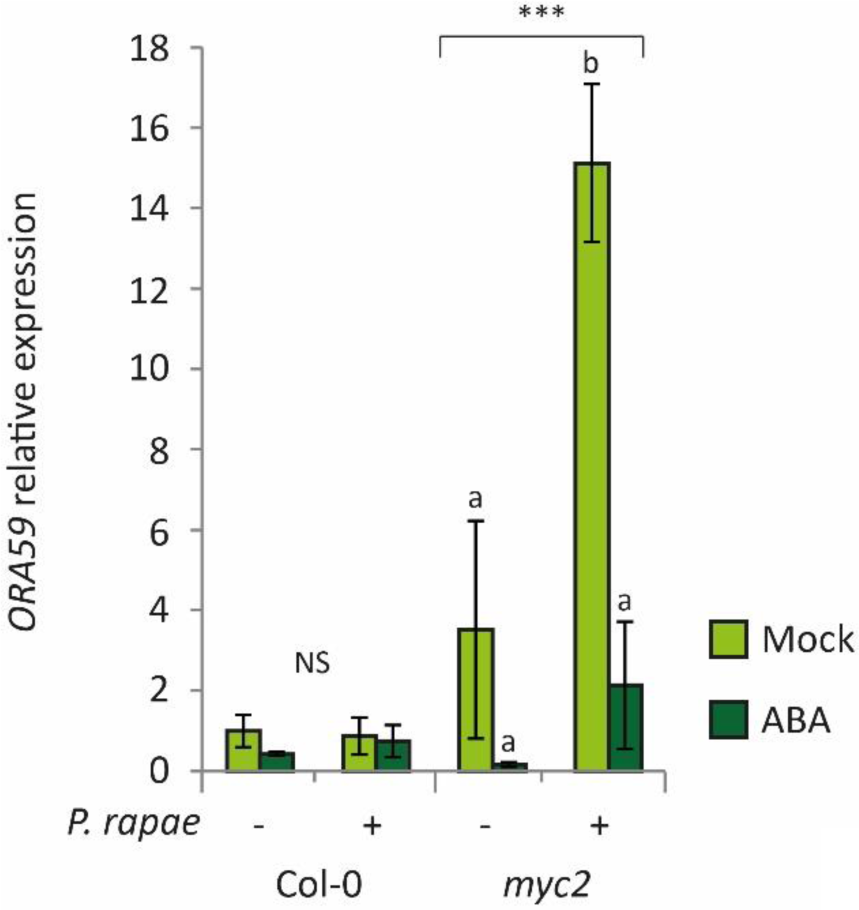
Effect of ABA treatment on *P. rapae*-induced *ORA59* expression. RT-qPCR analysis of *ORA59* gene expression at 24 h in leaves of Col-0 and *myc2* plants that were treated with a mock solution or with 100 µM ABA 24 h prior to infestation with *P. rapae*. Indicated are expression levels relative to mock-treated Col-0 plants at 24 h. Different letters indicate statistically significant differences between treatments of one line. Indications above the brackets specify whether there is an overall statistically significant difference between *myc2* and Col-0 (two-way ANOVA (treatment x genotype), LSD test for multiple comparisons; *** = *P*<0.001). Error bars represent SE, *n*=3 plants.

